# GAGER: gene regulatory network assisted gene expression restoration

**DOI:** 10.1101/2024.11.27.625595

**Authors:** Md Zarzees Uddin Shah Chowdhury, Sumaiya Sultana Any, Md. Abul Hasan Samee, Atif Rahman

**Author notes:** Corresponding author(s). E-mail(s); Contributing authors.

## Abstract

Gene regulatory networks are crucial for cellular function, and disruptions in transcription factor (TF) regulation often lead to diseases. However, identifying TFs to transition a source cell state to a desired target state remains challenging. We present a method to identify key TFs whose perturbation can restore gene expressions in a source state to target levels. Its effectiveness is demonstrated on datasets from yeast TF knockouts, cardiomyocytes from hypoplastic left heart syndrome patients, and mouse models of neurodegeneration. The method accurately identifies knocked-out TFs in the yeast dataset. In the cardiomyocyte dataset, it pinpoints TFs that, though not differentially expressed in many cases, exert significant regulatory influence on downstream differentially expressed genes. Finally, in the mouse model dataset, it identifies disease stage-specific TFs, improving similarity between healthy and diseased states at various time points. Unlike traditional approaches relying on differential expression analysis, our method uses network-based prioritization for more targeted and biologically relevant TF selection. These findings highlight its potential as a therapeutic tool for precise TF targeting to normalize gene expressions in diseased states.

## 1 Introduction

Transcriptional perturbation represents a promising avenue for molecular therapy. The advent of single-cell transcriptomics (scRNA-seq) has enabled significant advances in delineating pathological and physiological cell states. However, the identification of target genes for activation or repression to effectively transform a source cell state into a desired target cell state remains a largely unresolved challenge. This manuscript addresses this critical issue.

Accurate identification of transcriptional perturbation targets requires deep knowledge about the gene regulatory networks (GRNs) in both source and target cell states. Gene Regulatory Networks (GRNs) are very useful to describe many of the complex processes that underlie gene expression and regulation in a large number of biological systems. These networks define the complex relationships between genes, transcription factors, and other molecular entities that regulate and determine the behavior of cellular systems [1, 2]. In a GRN, nodes generally represent genes or transcription factors (TFs), whereas edges describe regulatory interactions between them [3].

In recent decades, substantial advances in the reconstruction of GRNs have been driven by large-scale datasets from technologies such as transcriptomics, epigenomics, and proteomics [4–8], alongside the development of sophisticated algorithms that infer these networks from high-dimensional data [9–14]. These GRNs are extremely important for biological research since they provide an approach to understanding the mechanisms of regulation that underlie biological processes, like cellular differentiation [15–17], gene knockdown effects [18, 19], and drug target prediction in diseases [20, 21]. Researchers simulate the response of the network to various perturbations and predict the outcome of genetic modification, which is of high value in therapeutic applications [13, 22].

One of the key applications for GRNs is in differential comparisons across different biological states, such as disease versus healthy tissues [23, 24], different developmental stages [25], or knockout and/or mutant models versus wild type [26]. Such differential studies will enable the identification of the genes that may be disease- or time-point-specific and therefore could be drug target candidates or uncover missing regulatory elements. However, in many approaches, the focus lies on sub-networks of smaller order—graphlet-based analysis [27], gene co-expression networks [23], or GRNs constructed from literature knowledge [24]. While these methods provide valuable insights, they come with challenges in scalability and interpretability.

Most importantly, most of these current approaches infer GRNs or regulatory relationships from scratch using different modalities of data. However, recent developments have enabled us to design highly accurate algorithms that infer GRNs from a number of data modalities, such as single-cell sequencing [10, 11] or bulk sequencing [28], motif analysis [29], ChIP-seq [30], proteomics, and epigenomics. These tools allow for the inference of whole GRNs from existing datasets, eliminating the need to develop separate methods for GRN reconstruction and subsequent differential analysis. This advancement simplifies the analysis and improves scalability, enabling researchers to focus on differential comparisons rather than the initial reconstruction of networks.

Despite these advancements, a significant limitation of existing methods is their inability to leverage GRNs effectively for guiding interventions that modulate gene expression in pathological tissues to mimic healthy tissues. To address the limitation of existing methods in effectively leveraging gene regulatory networks (GRNs) for therapeutic interventions, we have developed a novel algorithm named GAGER (GRN Assisted Gene Expression Restoration). This algorithm is designed to compare GRNs under two different conditions and identify specific genes whose manipulation could shift gene expression from a source condition, such as diseased tissue, towards a target condition, such as healthy tissue.

The core functionality of GAGER involves a forward simulation method that applies a series of perturbations to facilitate the transformation of a source (e.g., pathological) cell state into a target (e.g., physiological) cell state. This approach enables counterfactual inferences regarding how gene expression in the source state can be modulated to closely approximate that of the target state. By employing this method, we can hypothesize and test the impact of targeted interventions on gene expression restoration.

GAGER focuses on identifying differential regulatory edges between source (e.g., pathological) and target (e.g., physiological) cell states. This analysis is pivotal for pinpointing TFs that are central to the transcriptional regulation differences observed between the two states. By prioritizing TFs whose modulation could restore downstream gene expression, we aim to facilitate the transition of the source cell state towards the target cell state. In contrast to CellOracle [13], which can simulate the effect of perturbation or knockout of a TF given a list of TFs, our approach can generate a set of TFs to restore gene expressions to levels of a desired state.

We provide evidence of GAGER’s efficacy in three distinct application scenarios: 1) datasets from Saccharomyces cerevisiae under various TF knockout conditions [31, 32], 2) heart disease patients with Hypoplastic Left Heart Syndrome (HLHS) datasets [33], and 3) neurodegeneration mouse model data [34]. In these cases, our approach offered a highly focused list of genes for perturbation, facilitating improved alignment between source and target states upon restoring their expression.

The results from our study demonstrated improvements in cell state similarity and the identification of key transcription factors with substantial influence, proving that even a simple linear regression-based model can yield meaningful insights into complex biological systems. Our counterfactual approach differs from traditional methods that typically revolve around selecting differentially expressed genes (DEGs) or manually perturbing some TFs, followed by observing changes in gene expression [13]. Instead, our method focuses on targeted interventions, guided by prior knowledge of critical TFs, to more effectively restore gene expression patterns. This targeted strategy not only enhances the precision of our interventions but also highlights the simplicity and effectiveness of using a graph based and linear regression based method. Furthermore, our approach offers scalability and practical advantages by concentrating on fewer key genes, differentiating it from other large-scale analyses that often involve extensive datasets and numerous DEGs. These factors underscore the potential of our baseline counterfactual algorithm in GRN analysis for therapeutic interventions.

## 2 Results

### Analysis of gene regulatory networks to restore gene expression

We present a graph theoretical method to identify transcription factors (TFs) within a gene regulatory network (GRN) for perturbation to restore gene expressions. Our goal is to prioritize transcription factors (TFs) whose perturbation (up or downregulation) would make a source single-cell population similar to a target single-cell population in gene expression values. As an example of a practical use case, the source and target populations could be cells from a pathological and a control sample, respectively.

Our method, implemented in GAGER, is illustrated in Figure 1A. It takes as input two gene expression matrices from source and target cells. The first step in the method is to construct gene regulatory networks corresponding to the source and target cells i.e. the source GRN, *G*^source^ = (*V, E*^source^) and the target GRN, *G*^target^ = (*V, E*^target^) (details in Methods). GRNs are weighted networks where node weights denote the average expression values of the corresponding genes, and edge weights denote the strengths of regulatory effects. These regulatory edges are filtered based on the edge weights to retain only significant edges.

**Fig. 1:**
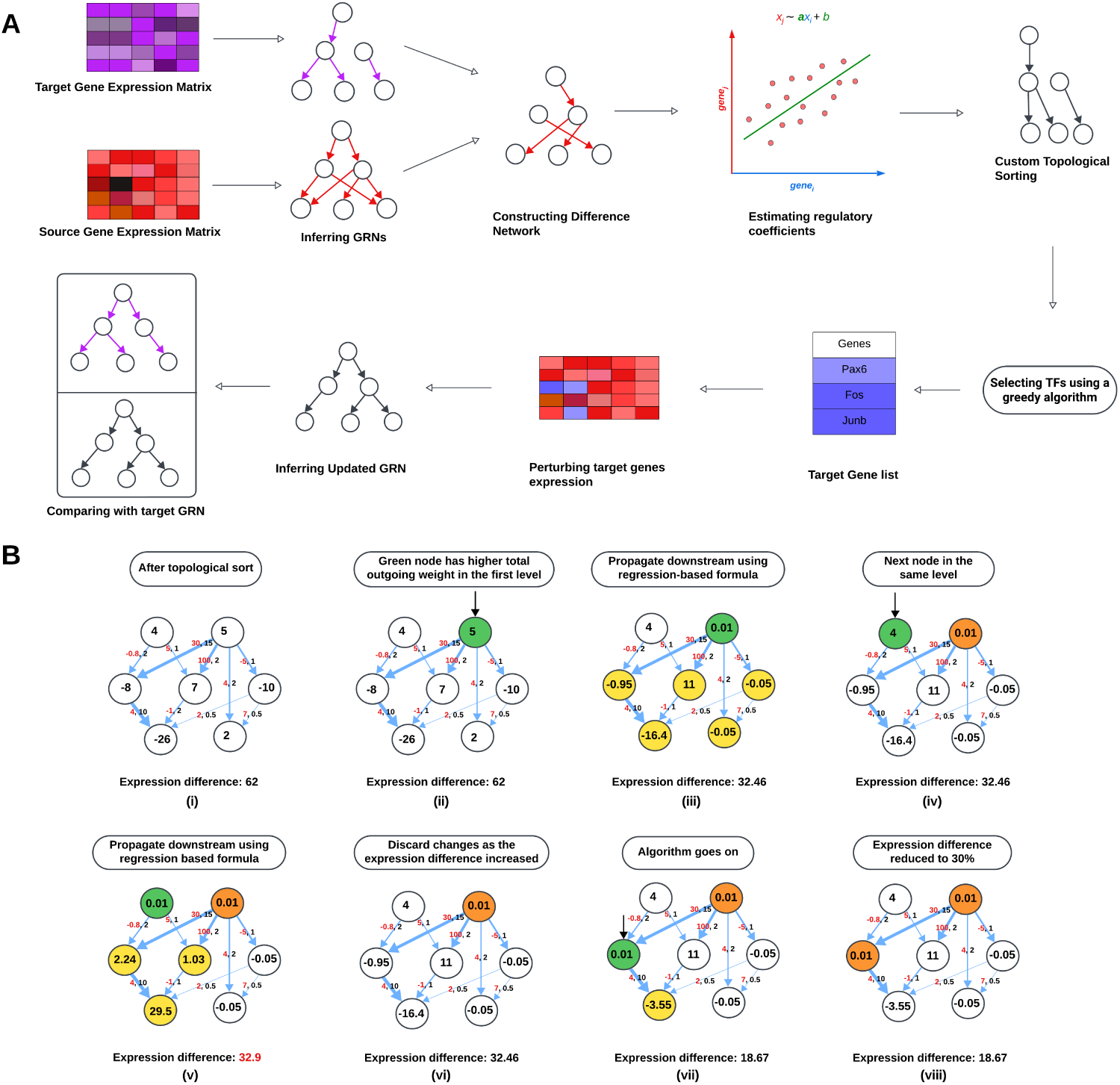
**A. Overview of GAGER.** The input consists of two gene expression matrices from source and target cells. GRNs corresponding to source and target conditions are inferred. A difference network is then constructed by identifying edges present in the source GRN but absent in the target GRN. Node values in this difference network represent the expression difference between the two states. To estimate how much the expression of a downstream node changes in response to perturbing a selected node, linear regression is performed. Next, nodes in this difference graph undergo a custom topological sorting. A greedy selection process is then used to choose candidate TFs for perturbation. The gene expression of the source condition for these selected TFs is adjusted to match the expression of the same TFs in the target condition. Based on this updated expression matrix, a new GRN is inferred. Finally, the updated source GRN and the target GRN are compared. **B. Greedy selection of TFs. (i).** The difference network after custom topological sorting, where nodes on the same level are prioritized based on their total weight of outgoing edges. Nodes are arranged level by level, with edges having higher weights shown by thicker arrows. Edges are labeled with regression coefficients (red) and regulatory weights (black). **(ii).** The TF node colored green is considered first due to its higher outgoing weight compared to the other node at the same level. **(iii).** Its expression difference is set to approximately 0, and the expression differences of the downstream nodes (yellow) are estimated using linear regression based formulas. This results in a decrease in the total absolute expression difference from 62 to 32.46, and so the TF is selected. **(iv).** The next candidate TF is highlighted in green. **(v) - (vi).** The expression differences of downstream nodes are estimated, but the total expression difference does not decrease, so expression differences of this node and its downstream nodes are restored. **(vii) - (viii).** The algorithm continues to explore nodes level by level, prioritizing nodes with higher out- going weight. If perturbing the TF expression difference reduces the overall difference, the node is selected, and updates are retained; otherwise, the node values are reverted. Eventually, only 2 nodes are selected (orange) and the total expression difference is reduced to 18.67. The algorithm may stop early if the total node expression difference falls below a predefined threshold; in this case, it is 30% of the initial total expression difference.

In the next step, these GRNs are compared to determine regulatory differences, and a difference graph or network *G*^diff^ = (*V, E*^source^ − *E*^target^) is created (see Methods). Here, regulatory differences refer to alterations in the regulatory interactions between genes across the two states, specifically focusing on edges present in the source GRN but absent in the target GRN. In addition, the difference network also encapsulates the differences in average gene expressions as node weights.

The prioritization of TFs first requires identifying all descendant nodes that may be influenced by changes in the expression of a selected TF. To achieve this, we perform a custom topological sorting of the nodes (see Methods for details). Topological sorting is an algorithm that orders the nodes in such a way that each node is placed before all of its descendant or downstream nodes. In addition, we prioritize nodes based on their total outgoing weights as they are likely to have more impact on expression of descendant nodes. Specifically, for two nodes *u* and *v*, if the sum of weights of outgoing edges from *u* are greater than that of *v*, then GAGER ranks *u* higher than *v*. Once the downstream nodes are identified, we perform linear regression to determine regulatory relationships. Linear regression allows us to estimate the regulatory coefficients, quantifying the effect of a selected TF’s expression on its downstream nodes. After identifying these relationships, we simulate perturbations to predict how changes in the expression of a TF will influence the expression of its downstream genes.

We then apply a greedy selection strategy to identify TFs for perturbation. An example of the algorithm is shown in Figure 1B and it is described in details in Methods. We iterate through the nodes according to the custom topological sort order. If the expression of a node in the diseased network deviates beyond a threshold (specifically, if the mean source expression of the gene is not within mean target expression ± 0.5 times the standard deviation of the target expression), the node is selected for perturbation simulation. We then set its expression difference close to zero. Then for each of its downstream nodes, a list of its parent nodes is obtained, and its change in expression is estimated by taking a weighted average of the expression differences of all its parents multiplied by their respective regression parameters according to the normalized weights of the connecting edges.

After updating the expression difference of a candidate node and propagating it to all its downstream nodes using the linear regression formulas, the new total expression difference in the difference graph is calculated by summing over the absolute differences. If this difference is less than the previous total expression difference, indicating a reduction in overall discrepancy between the source and target networks, the candidate TF is added to the list of selected TFs, and downstream node expressions are updated accordingly (see Figure 1B (ii) - (iii)). Conversely, if the total expression difference remains the same or increases, the previous expression differences are restored (see Figure 1 B (iv) - (v)). This iterative process continues until the total expression difference falls below a specified threshold which is empirically set to 30% of the initial total expression difference as default (see Figure 1 B (vi)- (vii)).

### GAGER identifies knocked out transcription factors in yeast datasets

First we assess the efficacy of GAGER in identifying knocked out transcription factors (TFs) using gene expression data. For this we use the dataset from [32] which contains scRNA-seq data from *Saccharomyces cerevisiae* samples: a wild-type control and 11 genotypes with 11 TF (GZF3, GLN3, GAT1, DAL80, DAL81, DAL82, GCN4, RTG1, RTG3, STP1, STP2) knockouts. The wild-type was grown in YPD and the knockouts were grown in various conditions including YPD. Finally, the knockouts were pooled together and sequenced along with the wild-type. The dataset is suitable for our experiment to see if our method can identify the knocked-out TFs as key regulators, since restoring their expression will revert the network to its original state.

To achieve this, we used the two datasets from this collection grown in YPD: one containing wild-type samples and another containing samples from the 11 TF knockout genotypes. Figure 2A shows the histogram of log fold change in gene expressions in the two datasets with those of the 11 knocked out TFs highlighted using vertical lines. We observe that the log fold changes lie within -0.5 and 0.5 except for the 11 outliers corresponding to the 11 knocked out TFs.

**Fig. 2:**
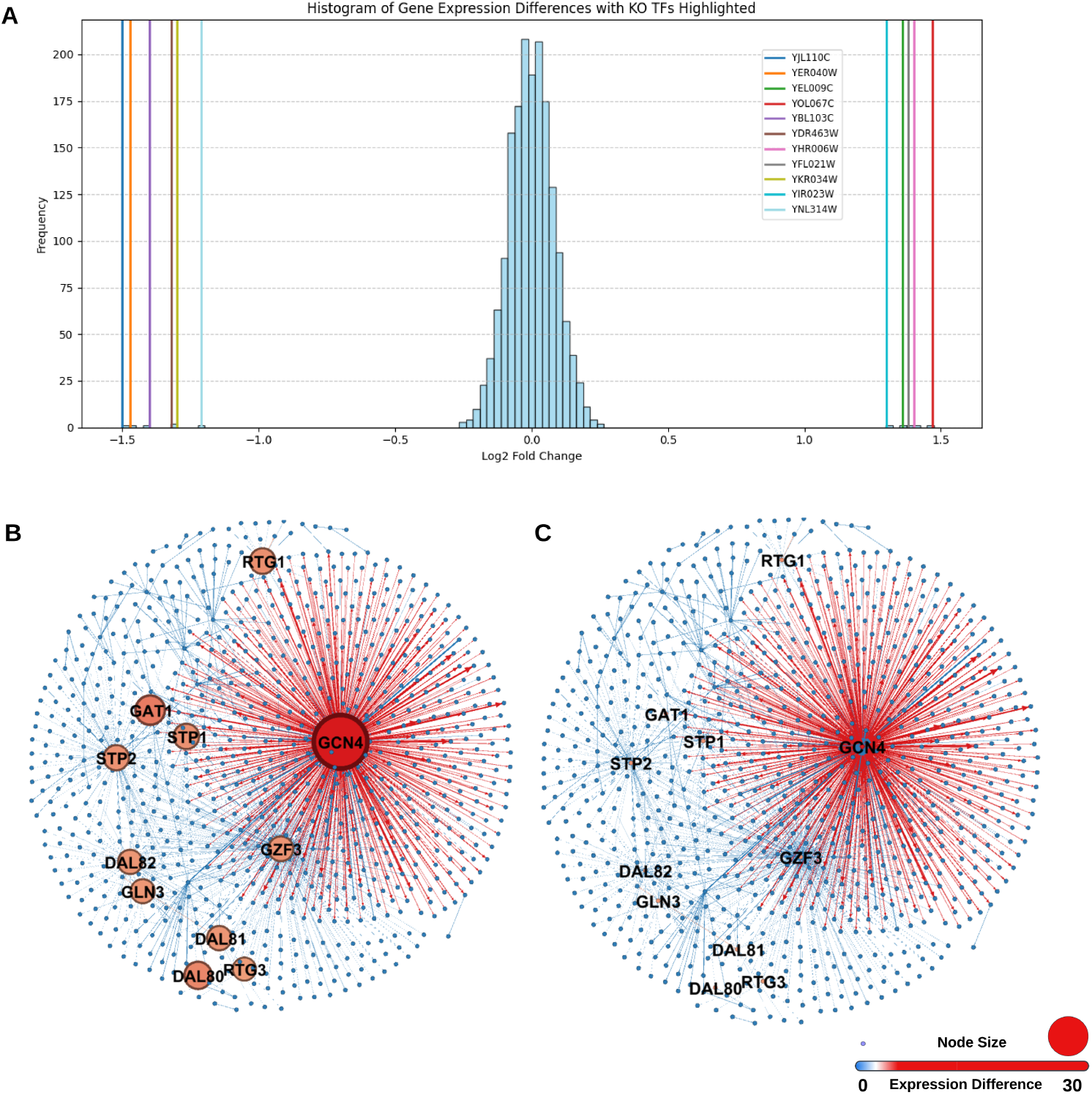
Analysis of the 11 knocked out TF *Saccharomyces cerevisiae* dataset. **A.** Histogram of log fold change in gene expression with the 11 outliers corresponding to the 11 knocked out TFs highlighted by vertical lines. **B.** Difference graph for the wildtype and 11 knocked out TF datasets. **C.** Difference graph after successfully identifying 11 TFs and seting their expression difference to 0.

We inferred the GRNs for both datasets and computed the difference graph based on expression differences. Figure 2B illustrates the difference graph for the 11 knocked-out TFs. We then applied our algorithm to identify TFs to perturb from this difference graph. Our method successfully selected the 11 TFs (GZF3, GLN3, GAT1, DAL80, DAL81, DAL82, GCN4, RTG1, RTG3, STP1, STP2), which are exactly the 11 knock-out genes. Figure 2C shows the difference graph after setting expression differences to zero and demonstrates that our method successfully selected these 11 TFs.

### GAGER finds regulators of significant transcription factors associated with hypoplastic left heart syndrome

Next we perform an experiment to contrast our approach with selecting transcription factors based on a simple differential expression test. To emphasize the difference between the two approaches, we use a dataset [33] containing induced pluripotent stem cell-derived cardiomyocytes (iPSC-CM) from 1 healthy control and 3 HLHS patients. Two patients were in group II who needed heart transplant or deceased. The group I patient survived transplant free and one was a healthy control subject.

We generated three difference networks (Control vs Group I, Control vs Group II, and Group I vs Group II) and run GAGER to identify TFs to perturb. We also find the differentially expressed genes (DEGs) using DESeq2 [35]. In Figures 3A-C, snapshots of the diffence networks are shown whereas Figure 3D summarizes the numbers of DEGs vs number of genes identified by GAGER in three conditions. The blue nodes are the selected TFs, and the red and yellow nodes are upregulated and downregulated genes, respectively.

**Fig. 3:**
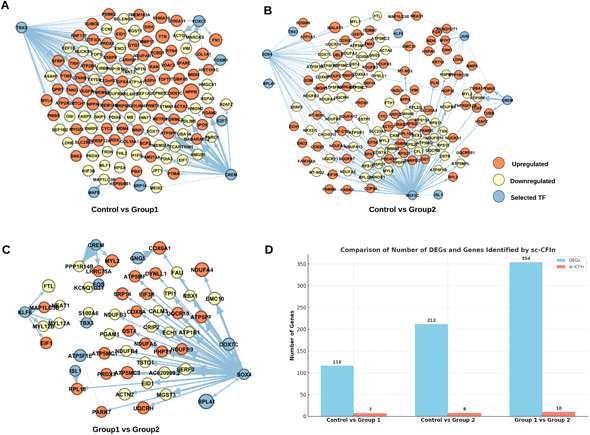
Difference networks showing selected transcription factors (TFs) and differentially expressed genes (DEGs) within 1 hop from the selected TFs in conditions related to hypoplastic left heart syndrome (HLHS), where the blue nodes are the selected TFs by our algorithm and the other nodes are DEGs, for **A.** Control vs HLHS Group 1 difference network, **B.** Control vs HLHS Group 2, and **C.** HLHS Group 1 vs Group 2. **D.** Counts of DEGs vs the number of identified genes by GAGER for different conditions.

In Control vs Group 1, the number of DEGs is 116 for adjusted P-value *<* 0.05 and log fold change *>* 0.5, but our method selected 7 genes (*CREM, TBX3, FOXC1, E2F7, MAFB, FOXM1, SRP14*). Similarly, for Control vs Group 2, the number of DEGs is 212, but our method identified 8 (*CREM, SOX4, KLF6, TBX3, MEF2C, ISL1, JUN, RPL41*). Finally, for Group 1 vs Group 2, the number of DEGs is 354, and our method detected 10 (*CREM, KLF6, SOX4, ISL1, TBX3, GNG5, COX7C, FOS, ATP5F1E, RPL41*).

We find that the numbers of TFs identified by our method are substantially lower than the corresponding numbers of DEGs. We also observe from Figures 3A-C that the TFs selected by GAGER have a large number of outgoing edges from them to the DEGs, and hence are regulators of those DEGs. Moreover, selected TFs are themselves not differentially expressed in a number of cases. This indicates that our method can effectively restore gene expressions by selecting only a few TFs which may not be DEGs but have a direct influence on DEGs.

### GAGER identifies disease stage-specific transcription factors to transition severe neurodegenerated mouse disease states to healthier states

One of our key goals was to analyze the dynamics of gene regulatory networks (GRNs) associated with disease progression. We examined a dataset [34] containing single-cell RNA sequencing data from 1685 individual microglia cells isolated from the hippocampus of mice with severe neurodegeneration and Alzheimer’s disease (AD)-like phenotypes, as well as control mice. This dataset includes samples from three to four CK-p25 mice and three CK control littermates at four time points: before p25 induction, 1 week, 2 weeks, and 6 weeks after p25 induction. The time-series gene expression matrices were suitable for generating GRNs week by week, enabling us to infer important transcription factors (TFs) using our algorithm and apply perturbations to observe how closely we could transition the diseased states to healthier states. After selecting TFs, we adjusted their expression to healthier levels, calculated the changes in their downstream genes according to the algorithm, and then used the updated gene expression matrix as input to SCENIC to infer the perturbed network. Our approach involved identifying candidate transcription factors (TFs) based on differential networks between healthy and unhealthy GRNs at various time points. The selected lists of genes corresponding to various stages are summarized in Figure 4A. These genes have significant roles in microglia development or neurodegeneration, as supported by existing literature.

**Fig. 4:**
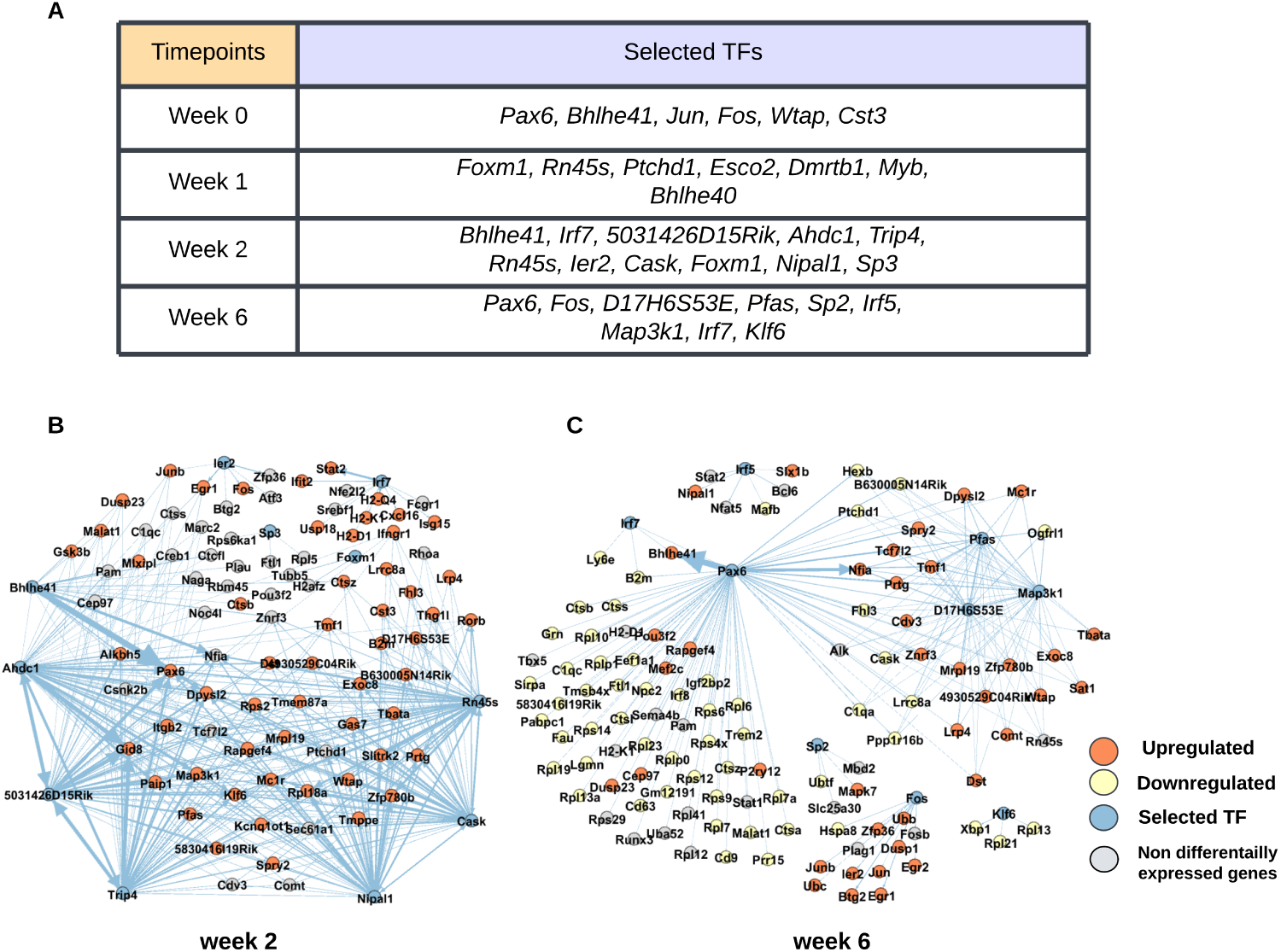
**A.** Selected TFs by our method at different time points. **B-C.** Two-hop neighborhood of the difference network from the selected transcription factors (TFs) between CK-p25 and CK mice **B.** at Week 2 and **C.** at Week 6. Blue nodes represent TFs selected by our algorithm, orange and yellow nodes represent upregulated and downregulated genes respectively, and other nodes are non-differentially expressed genes.

At Week 0, the candidate genes for perturbation included *Pax6*, *Bhlhe41*, *Jun*, *Fos*, *Wtap*, and *Cst3*. These genes are known for their critical roles in brain development or their activation in inflamed or injured microglia. For example, *Pax6* orchestrates neuronal development by ensuring unidirectionality and proper execution of the neurogenic program [36]. *Bhlhe40/41* are involved in regulating microglia and peripheral macrophage responses in Alzheimer’s disease and other lipid-associated disorders [37]. *Jun* and *Fos* are upregulated during microglia development, highlighting their roles as lineage-determining transcription factors (LDTFs) [38]. *Wtap* is activated in inflamed microglia [39], and *Cst3* maintains microglial homeostasis but is downregulated in neurodevelopmental disorders [40].

At Week 1, additional candidates included *Foxm1*, *Rn45s*, *Ptchd1*, *Esco2*, *Dmrtb1*, *Myb*, and *Bhlhe40*. *Esco2* ranked in the top 20 differentially expressed genes (DEGs) in microglia from an Alzheimer’s disease mouse model [41]. *Ptchd1* has been linked to neurodevelopmental processes, with insights provided by recent studies [42]. *Myb*, another candidate, is essential for definitive hematopoiesis [43].

At Week 2, the proposed TFs included *Bhlhe41*, *Irf7*, *5031426D15Rik*, *Ahdc1*, *Trip4*, *Rn45s*, *Ier2*, *Cask*, *Foxm1*, *Nipal1*, and *Sp3*. For instance, *Irf7* is crucial in modulating aberrant microglia activation states [44]. Research suggests *Cask* as a potential therapeutic target for central nervous system (CNS) disorders [45]. Additionally, *Sp3* has been implicated in regulating HIV-1 gene expression in human microglial cells through its interaction with COUP-TF (chicken ovalbumin upstream promoter) and *Sp1* [46].

At Week 6, selected genes included *Pax6*, *Fos*, *D17H6S53E*, *Pfas*, *Sp2*, *Irf5*, *Map3k1*, *Irf7*, and *Klf6*. *Sp2* regulates the cell cycle, and its deletion disrupts neurogenesis in the embryonic and postnatal brain [47]. *Irf5* has been linked to neuropathic pain by driving P2X4R+-reactive microglia [48], and targeting *Map3k1* alleviates inflammatory responses [49]. Moreover, *Klf6* is modulated by MiR-124 to influence microglia activation [50].

An important finding of [34], was the identification of two interferon response genes, *Ifitm3* and *Irf7*. Our results show that *Irf7* was consistently identified by the model during the final two weeks of the study.

These findings emphasize the effectiveness of our approach in identifying disease-stage-specific transcription factors (TFs) that play crucial roles in neurodevelopmental processes. Notably, some of these genes’ up- or downregulation is known to be associated with disease, and GAGER suggests a down- or upregulation, respectively, of those genes. For instance, *Jun*, *Fos*, *Irf5*, *Irf7*, *Esco2*, *Bhlhe40/41*, *Klf6*, and *Wtap* are upregulated in microglial neurodegeneration, and our method suggests downregulating them to bring their expression closer to healthy levels. Conversely, *Cst3*, which is downregulated, is suggested to be upregulated. These highlight the promise of targeted TF perturbations as a potential therapeutic strategy for addressing neurodegenerative diseases.

Figures 4B and C demonstrate that our selected TFs (blue nodes) have many edges towards differentially expressed genes (DEGs), indicated by orange and yellow nodes, underscoring their potential regulatory influence on these DEGs. Most of these two-hop neighborhood DEGs have important implications in microglial development and neurodegeneration, as evident from existing literature. For instance, qPCR analysis in mice previously revealed significant age-related microglial induction of MHC-I (Major Histocompatibility Complex) pathway genes, including *B2m*, *H2-D1*, and *H2-K1* [51]. We observe that at Weeks 2 and 6, *B2m* is regulated by *Rn45s* and *Pax6*, respectively, that have been selected by GAGER. At Week 2, *H2-D1* and *H2-K1* are regulated by *Irf7*, whereas at Week 6, both are regulated by *Pax6*. At Weeks 2 and 6, *Ctss* is regulated by *Ahdc1* and *Pax6*, respectively. Notably, *Ctss* plays a role in microglia-driven olfactory dysfunction [52]. *Mef2c* is another DEG regulated by *Pax6* at Week 6. In mice with microglial *Mef2C* deficiency, immune challenge leads to amplified microglial responses and negatively impacts mouse behavior [53]. Additionally, at Week 6, *Tcf7L2*, which plays a critical role in neurogenesis within the developing mouse neocortex [54], is downstream of *Pax6* and *Map3k1* selected by GAGER. Similarly, at Week 2, *Gsk3*. regulated by *Bhlhe41*, is known to control microglial migration, inflammatory responses, and neurotoxicity caused by inflammation [55]. These findings demonstrate our method’s capability to identify regulators of crucial DEGs.

## 3 Discussion

This study presented an approach for identifying transcription factors to perturb in order to transition a source cell state closer to a target state by restoring gene expressions. Current state-of-the-art methods include differential expression analysis [35], machine learning (ML) and deep learning (DL)-based perturbation frameworks [13] and hybrid approach including both algorithmic and experimental techniques [56].

We demonstrate that an algorithmic approach can yield meaningful insights into gene regulatory networks. We analyze an yeast TF knockout dataset and find that GAGER can identify knocked out TFs. Analysis of datasets from hypoplastic left heart syndrome patient datasets, and microglia from mouse models of neurodegeneration reveals that it can identify TFs that may not be DEGs but have direct influence on DEGs, and thus have important biological significance. While CellOracle [13] addresses related challenges, it primarily simulates single-step perturbations (e.g., TF knockouts), limiting its application to more complex scenarios. In contrast, GAGER incorporates multi-step perturbations, capturing the cascading regulatory effects throughout the network, which is essential for addressing intricate GRN changes. The TF list generated by GAGER may be provided to CellOracle for further analysis.

In future, our method can be extended in a number of ways. First, at present we use a specific tool SCENIC [10] for GRN inference. A future direction will be to explore other GRN inference tools [12, 13, 57, 58]. Second, due to the limited amount of data available to estimate the effect of changing expression of a gene on its downstream genes, we assume simple linear relationships In future, machine learning or deep learning algorithms [13, 59] may be explored. Finally, this approach may be tried on other omics data such as protein-protein interaction networks (PPI) [60].

## 4 Methods

### 4.1 Inference of gene regulatory networks

The first step in GAGER is inference of source and target GRNs from scRNA-seq data. A number of tools are available for construction of GRNs from scRNA-seq datasets such as SCENIC [10], CellOracle [13], scMTNI [11]. While other tools may also be used for this step, we use SCENIC (single-cell regulatory network inference & clustering) to infer directed weighted gene regulatory networks. SCENIC is a computational method for inferring gene regulatory networks and identifying cell states from single-cell RNA-seq data [10]. Inputs to SCENIC are a gene expression matrix and a list of transcription factors (TFs), and the output is a table containing regulators, their potential target genes, and the importance weight of each regulatory relationship. SCENIC utilizes the R/Bioconductor package GENIE3 (Gene Network Inference with Ensemble of trees) which is a method for inferring GRNs based on variable selection with ensembles of regression trees [9]. GENIE3 derives weights for the TFs as a measure to determine TF importance for target gene expression, facilitating the identification of TF-target regulatory links.

We run SCENIC with default parameters to infer two GRNs corresponding to the source and target cells. We provide as inputs gene expression matrices obtained from scRNA-seq datasets from the source (diseased) and target (healthy) cells. We then construct a directed weighted source graph or network (we use the terms graph and network interchangeably) from the output of SCENIC where each node in the network represents a gene and the directed edges represent regulatory relationships between the nodes. In the graph *G*^source^ = (*V, E*^source^), *V* is the set of all genes in the two datasets. If gene *g_i_* is inferred to be a regulator of gene *g_j_* in the source cells, there is a directed edge (*g_i_, g_j_*) ∈ *E*^source^ which is associated with the importance weight *w*(*g_i_, g_j_*) indicating the strength of the regulatory interaction from *g_i_* to *g_j_*. Similarly, we construct a target graph (*G*^target^ = (*V, E*^target^)).

### 4.2 Construction of difference network

The next step is to construct a difference network. First, we filter out edges from source and target networks to retain only important regulatory relationships. GENIE3 produces an output table containing genes, potential regulators, and their ‘importance measure’ (IM), representing the TF’s strength of regulation. A higher IM indicates greater confidence in the regulatory relationship. We observe that the distribution of IM follows a half-normal distribution (see Supplementary Figure S1). We normalize the IM and retain only edges with a normalized IM exceeding 0.1. This threshold was chosen empirically such that many low confidence edges are removed and the graphs contain only crucial regulatory edges, thereby simplifying analysis.

Then we construct the difference network *G*^diff^ = (*V, E*^diff^) by identifying edges that are present in the source GRN but absent in the target GRN. That is *V* is the same set of genes as the source and target networks, and *E*^diff^ = {*e*|*e* ∈ *E*^source^ and *e* ∈*/E*^target^}. For each node (gene) in the difference network, we also compute the difference in average expressions between the source and target networks i.e. to each node *g_i_* we assign

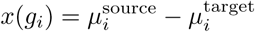

where *µ*^source^ and *µ*^target^ denote average expressions in the source and the target gene expression matrices for that gene, respectively.

Additionally, if the difference network contains cycles, the cycles are removed to ensure acyclic topology. To remove cycles, edges with the lowest importance in cycles are removed until the graph becomes acyclic.

### 4.3 Downstream node identification and estimation of regulatory coefficients

The process of identifying potential TFs for perturbation is based on whether the selected TF can reduce the total expression difference in the difference network. However, changes in the expression of a node may affect expressions of other genes. *Downstream nodes* of a selected node are those that have a direct path from the selected node in the source graph *G*^source^. These downstream nodes represent the genes whose expression may be affected by changes in the expression of the given node. We use a recursive algorithm to locate all downstream nodes of each node.

We also need to estimate the amount by which the expression of a downstream node will change in response to the perturbation of a selected node since the IMs do not have any statistical meaning. To model these relationships, we assume a linear relationship between the expression of a chosen node and its downstream nodes. That is we assume that the expression of a gene *x_i_* is related to the expression of a downstream gene *x_j_* as follows:

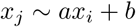

This choice of a linear relationship is motivated by the fact that we usually only have two datasets in the absence of replicates and the presence of dropouts in scRNA-seq datasets makes it difficult to utilise data at cell level. Moreover, linear relationship is known provide a good approximation of the behavior of the entire system [61].

We estimate *a* and *b* from average gene expression data in the source and target datasets i.e. *X_i_* = [*µ*^source^*, µ*^target^] and *X_j_* = [*µ*^source^*, µ*^target^]. If we have replicates or gene expression data at different time points, we use all the data

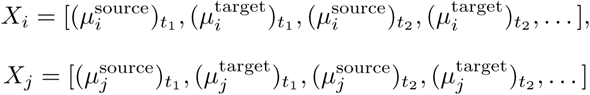

to estimate the parameters where (*µ*^source^)*_t_* denotes the average expression of the *i*-th gene in the source network at time or replicate *t*_1_. Then the difference in expression can be estimated using Δ*x_j_* = *a*Δ*x_i_*.

For each downstream node, a list of its parent nodes is obtained. Then, we apply the linear regression formula on the expression of each parent node and the downstream node to determine the regression parameters *a* and *b*. Next, the expression difference of the downstream node is estimated by taking a weighted average of the expression differences of all its parents multiplied by their respective regression parameters where the weights are the normalized importance scores of the edges connecting them. Here large values for weights correspond to actual regulatory interactions. Therefore,

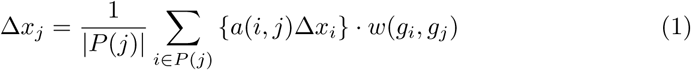

where *P* (*j*) = {*i*|(*g_i_, g_j_*) ∈ *E*^source^} is the set of parents of node *j* and *a*(*i, j*) is a regression coefficient obtained from the linear regression of *x_j_* on *x_i_*.

### 4.4 Custom topological sorting of nodes

We then topologically sort the nodes in the difference network. A topological sort arranges nodes in a directed acyclic graph in such a way that edges point from earlier to later nodes, reflecting network dependency relationships. Here we perform a custom topological sort which is described below:

- **Topological sorting**: Initially, a basic topological sorting algorithm is applied to the directed acyclic graph representing the difference network.
- **Layer assignment**: Next, nodes are assigned to layers. If a node *g_i_* does not have any parent, it is assigned to layer 0 i.e. *l*(*g_i_*) = 0. Otherwise, the maximum layer number of its parents is identified and it is assigned to the next layer i.e.

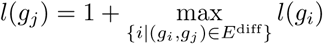

- **Custom sorting key calculation**: Then a custom sorting key is calculated for each node based on its total outgoing importance. The custom sorting key *I*(*g_i_*) is defined as

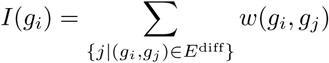

- **Updating topologically sorted node list**: The topologically sorted node list is updated layer by layer. Nodes are sorted in ascending order of their layer numbers and then each layer is sorted independently according to the custom sorting key. That is, if for two genes *g_i_* and *g_j_* in the same layer, *I*(*g_i_*) *> I*(*g_j_*), then *g_i_* is placed ahead of *g_j_*.

### 4.5 Selection of transcription factors to perturb

Finally, we select the transcription factors to perturb using a greedy algorithm. We consider the genes according to the custom topological sorting order. The order reflects the dependency relationships within the regulatory network and therefore ensures that a perturbation choice later will not have any effect on a node considered earlier. In addition, the order within each layer makes sure that nodes within each layer are positioned according to their likely impact on the network.

We consider the nodes one by one and if the mean source expression of the gene *µ*^source^ is not within *µ*^target^ ± 0.5*σ*^target^, it is a candidate TF for perturbation. Here, denote the mean target gene expression and the corresponding standard deviation, respectively. We then set its expression difference in the difference graph to zero. In practice, we set it to a small value (10^−5^) since setting it exactly zero may lead to numerical instability.

Upon selecting a TF node, we estimate the expression of downstream nodes in the source network using the linear regression formula mentioned earlier. The cumulative expression difference between source and target networks is calculated by summing node values in the difference graph. If modifying the current TF node’s expression difference contributes to an overall reduction in cumulative expression difference, it is retained as a candidate, and identification of additional TFs continues. However, if the total expression difference remains the same or increases, the candidate TF is discarded and expression difference for all nodes in the difference graph is restored. An example of the greedy TF selection process is illustrated in Figure 1B.

### 4.6 Iterative refinement

The process continues until the total expression difference falls below a specified threshold. We set it to 30% of the initial total expression difference as default. Selecting a lower value may result in perturbing a large number of nodes, which may not be feasible from a biological perspective. However, users can adjust this threshold according to their preferences. When the threshold is reached, the selected TFs are considered as the desired TFs for perturbation. We then change the expression value of the selected TFs accordingly in the source gene expression matrix and run SCENIC again by giving the perturbed source matrix as input to measure how much similarity has been achieved. When restoring expression of a gene in the source matrix, if the cells are the same in both matrices, we just restore the expression value in each cell from the target matrix into the source matrix. But if the sets of cells are not the same, we impute the expression values in the source matrix by fitting distributions for expressions in the target using KDE (kernel density estimation) and sampling from them. We measure the similarity in terms of the number of common edges in the perturbed source and target networks considering the edges with normalized importance greater than 0.1. Empirically, we find that a reduction in total expression difference to 30% of the initial total expression difference leads to more than 50% common edges in the source and target networks. In case expected result is not achieved, users may adjust the threshold and rerun the process.

### 4.7 Datasets

We analyzed the following datasets to obtain the results in this paper:

1. ***Saccharomyces cerevisiae Dataset*** (GSE125612): In this dataset [32], 12 yeast genotypes were utilized, comprising a wild-type control and 11 transcription factor deletions (GZF3, GLN3, GAT1, DAL80, DAL81, DAL82, GCN4, RTG1, RTG3, STP1, STP2). These strains were exposed to different environmental conditions and then the different strains were pooled together and sequenced. We used the mixed sample grown in YPD (GSM3564448). Additionally, untreated wild-type cells of genotype FY4/FY5 were included in a separate sample, grown under the condition YPD. This wild-type sample, represented by (GSM4039308), is part of the GSE125162 series and was used as the target.
2. **Hypoplastic Left Heart Syndrome (HLHS) Dataset** (GSE146341): The dataset [33] contains single-cell RNA sequencing (scRNA-seq) analysis of induced pluripotent stem cell-derived cardiomyocytes (iPSC-CM) obtained from patients with HLHS and healthy controls. scRNA-seq was performed on iPSC-CM from two group II patients - patient 7042 with heart transplant at 11 months and patient 7052 deceased at 2 months, a group I patient (patient 7464 surviving transplant free at 7 years of age), and a healthy control subject (1053).
3. **Progression of Neurodegeneration in Mouse Model** (GSE103334): This dataset [34] presents scRNA-seq data from 1685 individual microglia cells isolated from the hippocampus of mice with severe neurodegeneration and Alzheimer’s disease (AD)-like phenotypes, as well as data from control mice. It includes samples from three to four CK-p25 mice and three CK control littermates at four time points: before p25 induction, 1 week, 2 weeks, and 6 weeks after p25 induction (abbreviated as 0wk, 1wk, 2wk, and 6wk, respectively).

The CK-p25 mice, characterized by a genetic modification inducing the expression of p25 under the CamKII promoter, serve as a model for AD-like pathology and neurodegeneration. In contrast, CK control mice, unaltered genetically, provide a baseline comparison group devoid of specific alterations related to Alzheimer’s disease or neurodegeneration. This dataset offers valuable insights into the transcriptional dynamics of microglial cells throughout the progression of neurodegeneration in the CK-p25 mouse model.

## Declarations

### Data availability

The datasets and computer code produced in this study are available in the following databases:

- GAGER: code Github (https://github.com/PrinceZarzees/GAGER)
- *Saccharomyces cerevisiae* RNA-seq data: gene expression Omnibus GSM3564448 (https://www.ncbi.nlm.nih.gov/geo/query/acc.cgi?acc=GSM3564448) Omnibus GSM4039308 (https://www.ncbi.nlm.nih.gov/geo/query/acc.cgi?acc=GSM4039308)
- Hypoplastic Left Heart Syndrome (HLHS) RNA-seq data: gene expression Omnibus GSE146341 (https://www.ncbi.nlm.nih.gov/geo/query/acc.cgi?acc=GSE146341)
- Progression of Neurodegeneration in Mouse Model RNA-seq data: gene expression Omnibus GSE103334 (https://www.ncbi.nlm.nih.gov/geo/query/acc.cgi?acc=GSE103334)

